# Optimal Vaccination of a General Population Network via Genetic Algorithms

**DOI:** 10.1101/227116

**Authors:** Lloyd Sanders, Olivia Woolley-Meza

## Abstract

Herein we extend the work from Patel et al. (1) to find the approximate, optimal distribution of vaccinations of a virus spreading on a network with the use of Genetic Algorithms (GAs). We situate our investigation in an online social network, a Facebook graph of ~4000 nodes. Within this framework we investigate the performance of an optimized vaccine distribution scheme against that of a well known heuristic scheme: the vaccination of highly ranked nodes. We include also a baseline scheme of vaccinating random nodes. We show the algorithm is superior to rank scheme in low vaccine coverages, and performs comparably for most other coverage values, lending support to the optimality of this heuristic measure.

## Introduction

The role of vaccination in public health can hardly be understated. The reduction in rates of Polio and Measles are a direct result of the preventative effectiveness of vaccines. The ability of such prophylactics to save lives, and to a lesser extent reduce the financial burden on a society, is therefore of great importance to the scientific community.

The topic of vaccination is a large and complex field (2). There has been a fair amount of work done on the optimal method to vaccinate a population under various schemes. From the perspective of mean-field models, these schemes seek the best distribution over time, and generally with respect to some cost of the vaccine. Under these deterministic models one can employ methods from Optimal Control to find analytical solutions to the problem (3–5), for example: vaccination in a two-strain Tuberculosis model (6). This forward and back optimization is computationally, highly effective. But the lack of flexibility of the methodology works against it.

Mathematical Epidemiology has increasingly turned to modeling of spreading processes on complex networks (7) as they relax some of the harder assumptions of mean-field models by providing a contact structure. With this added complexity comes rich, realistic dynamics (Fig. 1), but the models often move further from analytical tractability. Although, some aspects of epidemics on networks can be characterized analytically, such as the critical translatability needed to infect a sizable fraction of the network (akin to the basic reproduction number), and spreading rates (2). As these models grow in sophistication, so do the vaccination schemes.

**Fig. 1.**
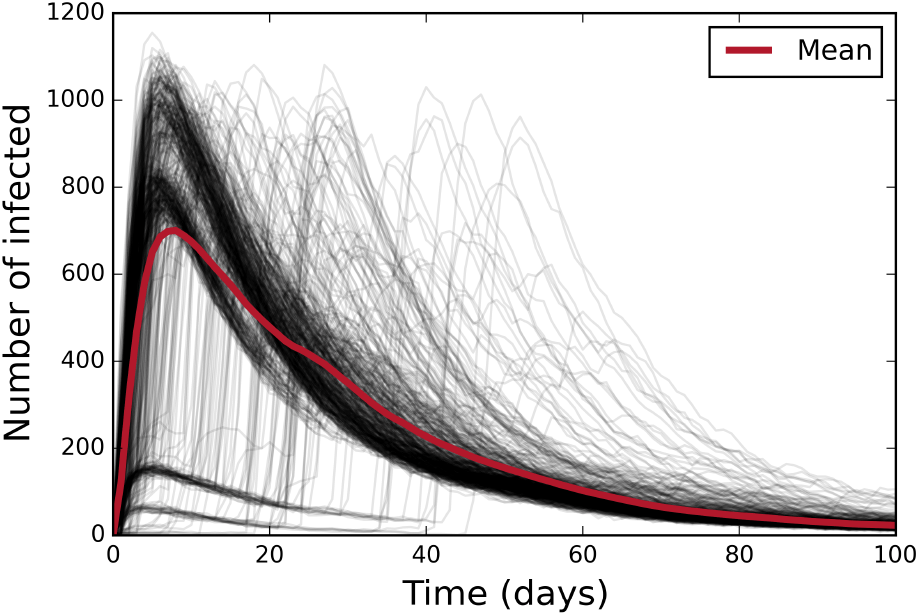
Here we model simple SIR contagion, Eq. (1), on the Facebook network (Fig. 2). We simulate 500 outbreaks (with random starting locations - a single node infected each time) shown individually as the grey lines, whose mean is given by the red. Recovery time is modeled after Influenza, of six days (*γ* = 1/6), whereas, *R*_o_ = 2.5, so we set *β* = γ **•** *R*_o_ ≈ 0.42.

Vaccination schemes on networks can readily adopt the frame work of Statistical Physics and map it to a site percolation problem – hence some characterizations, mentioned above, are possible. Given this thread of thinking, some schemes hope to increase the percolation threshold (by removing nodes – vaccination) and, thereby reduce the spread of the virus. Alternately, many schemes are based on heuristic models characterized by node measures of the network, such as degree correlations, or differing types of centrality measures, such as: betweeness, eigenvector, random-walk, closeness (2, 8). All of these methods assume global knowledge of the networks, which in reality is not often available. There has been work on vaccination methods on networks, with only local properties, such as acquaintance vaccination (2). Finally there are other more specific targeting schemes such as ring vaccination (9).

Although this subfield is well populated, we believe there is a need for method to find an (approximate) optimal solution, for use in the field, and also to benchmark these heuristic measures mentioned. We therefore look to the work of Patel et al. (1), to extend their work to networks. Patel et al. devised a scheme to deliver the optimal amount of vaccinations to each age group in their age-stratified, metapopulation Influenza model. They considered two numerical optimization schemes to benchmark against a random distribution of vaccines; namely Genetic Algorithms and Random Hill Climbing with mutation. The former was found to be optimal, and hence is the focus of this study. Genetic Algorithms, succinctly describe by Hamblin (10), “*…are a heuristic global optimization technique mimicking the action of natural selection to solve hard optimization problems…*” The work of Patel and co. was novel, but lacked generality to a network contact structure, as opposed to a metapopulation model. It is here where our work sits. We expound upon their model to that of a general contact structure network and at the same time bring parsimony to certain aspects of the algorithm to create a general, flexible method to find an optimal strategy for vaccination on general networks.

## Spreading and Vaccination

For our work, we analyze a simple contagion model; we use the SIR model as our basis, for its generality with respect to mimicking many viral traits. We adapt the model to a network, where each node on a network is a person. We assume that people can be infected by infected nearest neighbors with a probability

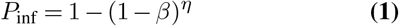

where *η* is the number of infected nearest neighbors, and *β* is the probability that one infected neighbor with infect another neighbor in a given timestep *dt*. Nodes recover with a probability, *γ*, in a given timestep *dt*. These surprisingly rich dynamics are shown in Fig. 1.

In our model, nodes which are initially Susceptible can be vaccinated. That is, vaccination has the role of shifting the health state from Susceptible, to Recovered, and is assumed to be 100% effective. Immunity is considered lifelong.

### Optimization Scheme

For the optimization scheme, we follow the path laid out by Patel and co. (1), for which we will briefly go over here, with an adjustment appropriate for networks.

In their work, their spreading model consists of an age stratified, metapopulation model. Here we remove the stratification (such that all nodes are homogeneous) but increase the contact structure to a network of *N* nodes (people). We suppose that we have *n_v_* vaccines available (one per person, assuming efficacy of the vaccines to be 100%) at the beginning of the simulation (sans infection) that we can distribute. Here, we conduct the model as follows.

Construct *m* individuals, conceptually thought of as vaccination strategies, which are vectors of genes, where each vector is the genome of the individual (in the nomenclature of GAs). We denote the *i^th^* individual/strategy as 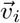. Each index of an individual maps to a distinct node on the network. The elemental value on that index, referred to as the gene, or locus, can be 0 or 1, denoting that the node to which it maps is either non-vaccinated or vaccinated, respectively. Note then that 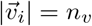. An outbreak is initiated post-vaccination, where one random node is infected per simulation. For each individual, the simulation of a virus spreading on the network is run through *e_n_* times (ensemble size). The number of recovered nodes (excluding those initially vaccinated) are summed to give the total total number of nodes affected by the disease. The lower this number, the higher the so-called fitness of the individual. We encode this mathematically as,

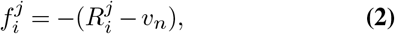

where 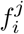 is the fitness of the *i*^th^ individual, for the *j^th^* simulation out of the ensemble, where 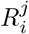 denotes the final number of recovered at the end of the simulation. Therefore, the mean fitness, for a given individual is

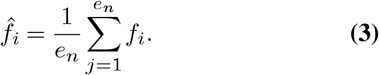

Note: Simulations are conducted until no infected nodes are left.

For each individual, the mean fitness is found for a given generation. The whole procedure is iterated over many generations, *g*. Between each generation, genetic information exchange occurs between individuals. Following Patel et al.’s work, we use both Tournament selection (below) and elitist selection (10): select the top fittest half of the individuals to be immediately passed to the next generation. Tournament selection is used to construct the remaining individuals such that the total number of individuals per generation remains constant. The tournaments are as follows.

### Mating Scheme and Tournaments

To construct the remaining individuals to pass onto the next generations, firstly, select a pool of 10 individuals without replacement from the whole set. From these find the one with the highest average fitness: the paternal individual. Construct another tournament from the current pool of individuals sans the paternal individual, of the same size. Select the one with the highest fitness: the maternal individual.

From here we mate the two via crossover breeding. The mixing of gene states (0 or 1) is not necessarily 50/50. We designate this as the crossbreeding factor, *c_B_*(< 1).

In this situation, to construct the offspring individual from the paternal and maternal, we consider each gene index (the vector index) in sequence. With a probability *c_B_* we select the gene from the paternal, otherwise we select it from the maternal individual. Once all genes have been chosen, we need to balance the number of vaccines issued in the offspring individual; namely it may have greater or less than the total number of vaccines required, *υ_n_*. If there are a greater number of vaccines issued to the offspring, we collect all indexes which have a value equal to unity. Without replacement, we select, at random, the same number of genes as the excess number of vaccines issued. Those genes are therefore set to 0. In the case too few vaccines are issued, we collect all null genes. Without replacement, we select at random the same number of genes as needed to make up the total number of vaccines issued per individual. We set those genes selected to unity. This final adjustment on the offspring individual serves as a mutation step in the creation process. We here depart from the original authors’ scheme due to the differing nature of our model and in seeking a more parsimonious algorithm.

### Convergence

We set the convergence of the algorithm in a similar fashion to before: if the top mean fitness of an individual has not changed after *c* generations, or the algorithm has exceeded *g* generations of computation, we assume convergence.

### Rank Vaccination

To benchmark our work, we also use a commonly accepted heuristic strategy: the vaccination of nodes with high degree of nearest neighbors (11). To conduct this strategy, we decompose the network into nodes, rank (hence the namesake) in a descending fashion, in terms of degree. For a given percentage of vaccination coverage, we take that percentage of the list, and vaccinate those nodes, starting from the node with the highest degree/rank. As the vaccine coverage increases, we vaccinate more and more nodes with a lower and lower degree. Explicit implementation is left till below.

### Random Vaccination

To benchmark our work, we measure the approximate optimal vaccination scheme against a random distribution of vaccines over the network, irrespective of the nodal hierarchy. Again, we run the random distribution over an ensemble of simulations, and take the mean fitness to represent the benchmark vaccination proficiency level.

## Results

### Model Initialization and Parameters

To investigate the performance of our methodology we require a representative real-world contact network. We have thus chosen to analyze a Facebook network provided by SNAP (12). This network is undirected, with 4039 nodes, 88234 edges, and a diameter of 8 - a size which is not trivial, but still computationally tractable. This network was investigated in (13) and is displayed in Fig. 2.^1^

**Fig. 2.**
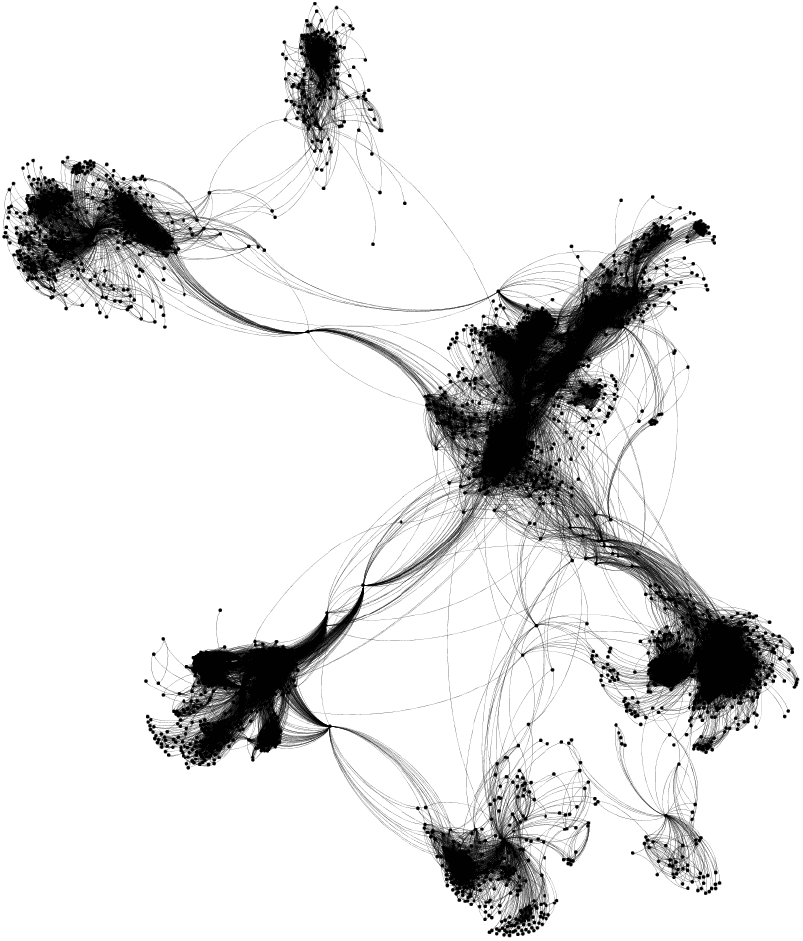
We base our calculations on an anonymized Facebook network provided by SNAP (12). The network has 4039 nodes, 88234 edges, and a diameter of 8. Illustrated by Gephi(14).

To compare the GA vaccination scheme to the Rank and random benchmark schemes, we consider the same network structure for each case.^2^ During the simulations, the viral parameters and simulation parameters are held constant. Namely, we set *γ* = 1/6, and *γ* = *β* · 2.5. All simulations are initially infected with a single node that varies randomly for each simulation, but this variation is accounted for by running the simulation many times (the ensemble size). We consider a range of vaccine coverages [5-90 %], and run the simulations on the network, such that for the given spreading process, the simulation is run until there are no more infected nodes on the network.

For the GA, we use 50 individuals, The initial (random) individuals for the GA (and the random scheme) are chosen prior to infecting the network for any simulation. For the Rank algorithm, we immunize according to the instructions set out in the previous section for this algorithm. The network is immunized before selecting the infected node. For the GA, the top 25 fittest individuals are passed to each new generation.

The remaining 25 are created via a tournament scheme composed of 10 individuals. We select our crossover breeding percentage to be 80%, as in Patel et al. We conduct the GA over 20 generations, or until the algorithm converges. The parameters for our simulations are found in Table 1.

**Table 1.** The parameters used to define the virus, and the vaccination are contained in top of the table. The bottom of the table is reserved for parameters of the GA.

| Parameter | Value(s) (Increment) |
| --- | --- |
| Recovery prob., $\gamma$ | 0.167 |
| Contagion prob., $\beta$ | 0.42 |
| Vaccination coverage, $v_n$ | [0.05, 0.9] (0.05) |
| Ensemble Size, $e_n$ | 250 |
| Generations, $g$ | 20 |
| Convergence Number, $c$ | 4 |
| Crossover Breeding Prob., $c_B$ | 0.8 |
| Tournament Size | 10 |

### Vaccination Scheme Performances

In Fig. 3, for each vaccine coverage value, we show the the mean fitness between all three schemes: GA, Rank, and the random benchmark (including no vaccination scheme at all).

**Fig. 3.**
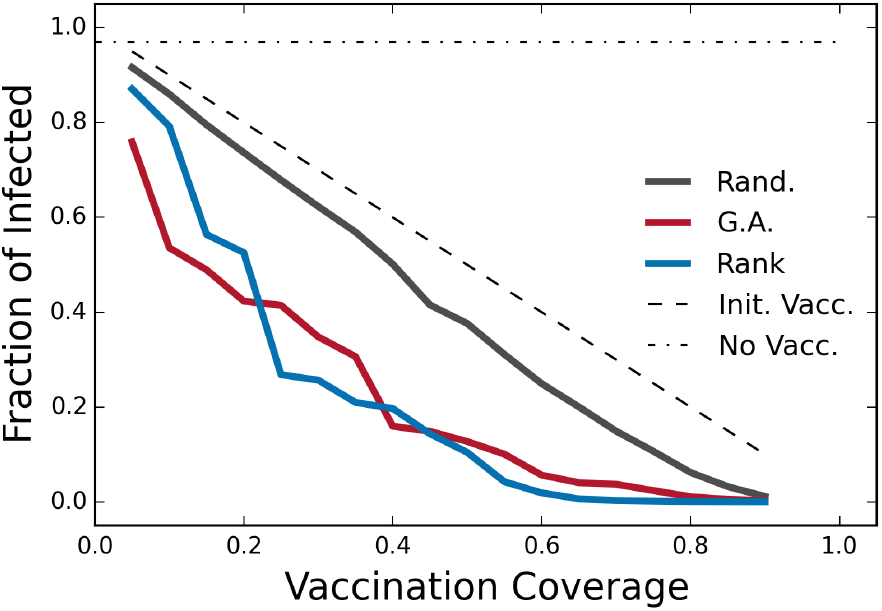
We show the performance of the GA approach (‘G.A.’, red) against the common high-degree strategy (’Rank’, in blue), and a random vaccine strategy (’Rand.’ in black). We show the worst-case scenario ‘Init. Vacc.’, where only those vaccinated are untouched by the disease. We also include no vaccination (’No Vacc.’ Black dot-dashed line). Simulations are conducted on the network shown in Fig. 2 for a simple contagion process [*β* = 1/6, *γ* = *β ·* 2.5, Eq. (1)]. On the y-axis we show the average number of people infected (and then recovered) from the virus, less the number of vaccines issued for each simulation. As expected, the mean infected of the disease drops linearly with a random strategy. But we see a marked difference in the effectiveness of the GA, and Rank schemes. These algorithms are able to produce far more optimal arrangements at all vaccine coverage values. Initially the the GA outperforms the Rank, but passed 20% coverage, the Rank algorithm performs better. It is entirely possibly that, given enough resources, the GA should perform as well, if not better than the Rank scheme. GA results are average over 250 simulations; Rank and random results over 500 simulations. The remaining parameters are housed in Table.

We see in Fig. 3 the comparison between the oft cited Rank strategy, and the GA. Initially, the GA is the most effective for low vaccine coverage values, after which it is eclipsed by Rank algorithm. One notes, that aside from the coverage values of [0.25 - 0.4], the GA closely mimics the efficacy of the Rank algorithm. We assume that given enough computational time, the GA, should find a strategy similar to the Rank algorithm in the [0.25 - 0.4] frame, and here is likely stuck in some local minimum.

The random strategy is approximately linear in its effectiveness, whereas, the GA and Rank responses could likely be described as quadratic. In comparison: To ensure that half of the population is safe from the virus, the random strategy on average requires ~40% coverage rate, whereas the GA strategy requires ~15% coverage - an extremely efficient response.

## Discussion and Conclusion

We have extended the work by Patel et al. (1) to consider a general network structure with a simple spreading process upon it. We have shown, via Fig. 3, that our Genetic Algorithm scheme was able to find an approximate, optimal vaccination strategy, for the network considered in Fig. 2, that consistently beats the random scheme for any value of vaccine coverage. Through this we show the value of this meta-heuristic approach: With the advent of cheap computational power, one can find the optimal vaccination scheme for highly dimensional, complex models. The flexibility of this model, coupled with its simplicity, is its strength.

One should be cautious on generalizing these results to all graph types (ER-random, scale-free, and small world), as we have only investigated a single graph to compare these two heuristic measures. It is likely that the GA heuristic, given enough computational time, should find an approximation to the optimum spread of vaccines across any graph structure. How well the Rank algorithm performs, is up for investigation. The GA framework built here, gives one the opportunity to investigate the Rank’s efficacy across these graph types.

### Future Work

As this work is a preliminary use-case investigation, there is much to do in terms of future work with respect to this framework. Simply, one could test the sensitivity of the results with respect to viral parameters, and also, obviously with respect to network structure. It could be, that the GA is more efficacious with respect to some structure classes of networks that others. If this is then the case, benchmarking this algorithms against other meta-heuristic optimization algorithms, such as Simulated Annealing (16), would be beneficial.

As this method is effectively rather simple, the cost of computation is non-trivial. One could investigate the computational cost of convergence, with respect to the algorithmic parameters, namely: tournament size, individual size, etc. Finally: It will be interesting to see how the methodology can be adapted to other optimization problems on networks, such as managing disaster spreading via external resources (8), or in situations of complex contagion, such as advertising and opinion spreading [which would likely be reflected through changes to the fitness/cost function, Eq. (3)].

### Heuristic strategies inspired by Gas

We believe that the GA scheme offers more than simply a method to find the optimal solution to the vaccination problem on networks: It can be used to inspire other heuristic strategies of vaccination. Namely, when conducting simulations, we believe there will likely be certain nodes of greater importance to minimizing spread through their own vaccination. It is therefore likely, that over a range of coverage values, these nodes are picked out via the algorithm more than others – essentially weighting their importance more. Comparing the likelihood of vaccination for a given coverage as a function of nodal degree (or any other node measure on a network) could then inspire, or help validate some of the heuristic schemes mentioned in the Introduction, which are based on these nodal properties.

### Machine Learning based on GA feature sets

Looking further ahead: The GA can also be used as a basis to create a training set for Machine Learning algorithms with respect to vaccination schemes. One could pose the question: Can we teach an algorithm, given a snapshot of a susceptible network, the viral parameters, and the vaccine coverage, the likely best nodes to vaccinate? We believe, the GA framework explicated here, could be up to the task. Let us outline a possible avenue of investigation.

One could generate many different synthetic networks (be they small world, lattice, random, or scale-free), along side real network datasets, to run large ensembles of outbreaks with differing viral parameters. In each case, we could use the GA to find the approximate optimal solution given the appropriate cost function.

Once the vaccination scheme is found, a feature set could be developed where the viral parameters, network, network statistics, degree distributions and vaccine coverage are included. The output could be the proportion of vaccines given to each nodal degree in the network degree distribution. This would then constitute as a data point in a training set. Given a large enough training set, and using the appropriate machine learning algorithms, one could then train the algorithm to give out the vaccination scheme, given the inputs above. In so doing, one would have developed a machine learning algorithm to vaccinate populations, given a snapshot of the network.

## ACKNOWLEDGEMENTS

The authors would like to thank Caleb Koch, and Kaj Kolja Kleineberg for their discussions on the manuscript.

This work was partially funded by the European Community’s H2020 Program under the funding scheme “FETPROACT-1-2014: Global Systems Science (GSS)”, grant agreement 641191 “CIMPLEX: Bringing CItizens, Models and Data together in Participatory, Interactive SociaL EXploratories” (http://www.cimplex-project.eu).

## Footnotes

^1^This network was constructed and anonymized from all users participating in a Facebook application. See (13) for more details.

^2^The network is constructed from the python package NetworkX (15).

## Bibliography

1. Rajan Patel, Ira M Longini, and M Elizabeth Halloran. Finding optimal vaccination strategies for pandemic influenza using genetic algorithms. Journal of Theoretical Biology, 234(2): 201–212, 2005.

2. Zhen Wang, Chris T Bauch, Samit Bhattacharyya, Alberto d’Onofrio, Piero Manfredi, Matjaž Perc, Nicola Perra, Marcel Salathé, and Dawei Zhao. Statistical physics of vaccination. Physics Reports, 664:1–113, 2016.

3. Maia Martcheva. Introduction to Mathematical Epidemiology, volume 61. Springer, 2015.

4. Oluwaseun Sharomi and Tufail Malik. Optimal control in epidemiology. Annals of Operations Research, 251(1-2):55–71, 2017.

5. Rachael Miller Neilan and Suzanne Lenhart. An introduction to optimal control with an application in disease modeling. In Modeling Paradigms and Analysis of Disease Trasmission Models, pages 67–82, 2010.

6. E Jung, Suzanne Lenhart, and Z Feng. Optimal control of treatments in a two-strain tuberculosis model. Discrete and Continuous Dynamical Systems Series B, 2(4):473–482, 2002.

7. Romualdo Pastor-Satorras, Claudio Castellano, Piet Van Mieghem, and Alessandro Vespignani. Epidemic processes in complex networks. Reviews of Modern Physics, 87(3):925, 2015.

8. Lubos Buzna, Karsten Peters, Hendrik Ammoser, Christian Kühnert, and Dirk Helbing. Efficient response to cascading disaster spreading. Physical Review E, 75(5):056107, 2007.

9. David Greenhalgh. Optimal control of an epidemic by ring vaccination. Stochastic Models, 2(3):339–363, 1986.

10. Steven Hamblin. On the practical usage of genetic algorithms in ecology and evolution. Methods in Ecology and Evolution, 4(2):184–194, 2013.

11. Romualdo Pastor-Satorras and Alessandro Vespignani. Immunization of complex networks. Physical Review E, 65(3):036104, 2002.

12. Jure Leskovec and Andrej Krevl. SNAP Datasets: Stanford large network dataset collection. http://snap.stanford.edu/data June 2014.

13. Jure Leskovec and Julian J Mcaule. Learning to discover social circles in ego networks. In Advances in Neural Information Processing Systems, pages 539–547, 2012.

14. Mathieu Bastian, Sebastien Heymann, Mathieu Jacomy, et al. Gephi: an open source software for exploring and manipulating networks. ICWSM, 8:361–362, 2009.

15. Aric A. Hagberg, Daniel A. Schult, and Pieter J. Swart. Exploring network structure, dynamics, and function using NetworkX. In Proceedings of the 7th Python in Science Conference (SciPy2008), pages 11–15, Pasadena, CA USA, August 2008.

16. William H Press. Numerical recipes 3rd edition: The Art of Scientific Computing. Cambridge University Press, 2007.

